# Gas extraction under intertidal mudflats is associated with declines in sediment grain size and minor changes in macrozoobenthic community composition

**DOI:** 10.1101/2023.05.09.539962

**Authors:** Paula de la Barra, Geert Aarts, Allert Bijleveld

## Abstract

1. In intertidal environments, land subsidence may change the local flooding regime and sediment composition, two main drivers of the macrozoobenthic community structure. In the Dutch Wadden Sea, a UNESCO world heritage site, gas extraction has resulted in an average subsidence of up to 2 mm y^-1^ of intertidal mudflats. These mudflats support a highly productive macrozoobenthic community, which offers important resources for birds and fishes. To what extent land subsidence due to gas extraction affects sediment and macrozoobenthos remains unknown and increasingly important given sea level rise.
2. Taking advantage of a monitoring program, we evaluated the effect of anthropogenically caused land subsidence on sediment composition and intertidal macrozoobenthos. Nearly 4600 points were sampled yearly (2008-2020) across the Dutch Wadden Sea, allowing us to compare sediment composition and macrozoobenthos biomass within and outside the subsidence area while controlling for the main drivers of these variables. We also compared population trends within and outside the subsidence area for 31 species with different habitat use in terms of depth and sediment composition.
3. Sediment median grain size decreased in the subsided area at 1 µm y^-1^ while on average remaining constant in other intertidal mudflats. Mud fraction was 3% higher within the subsided area throughout the studied period. This had no effect on the total biomass of macrozoobenthos. The biomass of species that use deeper areas increased within the subsidence area compared to outside, and the opposite was true for species using shallower places, but comparable patterns were also found in an area not affected by subsidence.
4. Changes in median grain size could be happening, and minor changes in macrozoobenthic community composition. For a successful implementation of the “hand on the tap” principle in the Wadden Sea, it is necessary to define beforehand the relevant variables that represent the natural values, implement proper monitoring, and define threshold values above which effects are not acceptable. We propose median grain size, mud fraction and macrozoobenthic biomass as good measures of the natural values of the Wadden Sea, and the methods used here as a way for identifying anthropogenic effects on them.

## Introduction

The extraction of underground oil or gas modifies the structure of subterranean layers and can lead to land subsidence with consequences for the environment at different scales (Milliman & Haq 1996, Ko & Day 2004). As a consequence, habitat availability for plants and animals is also affected, and local ecosystems can be transformed (e.g. Li et al. 2019, Turner & Mo 2021). When land subsidence occurs in intertidal areas – regions that are submerged during high tide and emerge during low tide-it alters the local flooding regime. If subsidence rates are low relative to the sediment supply by water, subsidence can be partially or completely compensated by increased sedimentation rates (Wang et al. 2018). In mudflats, the equilibrium mud content of the sediment primarily depends on sediment deposition fluxes (Colina Alonso et al. 2022), and therefore subsidence might affect the mud content of the affected area. At high subsidence rates, however, intertidal areas may almost permanently inundate. This could be aggravated by increased erosion, as deeper areas permit stronger waves and shear stress (Wang et al. 2018). Both sedimentation and erosion, together with sediment supply, will ultimately determine the sediment composition of a particular area in a mudflat. Because subsidence changes depth, which partially determines sedimentation and erosion regimes, subsidence may ultimately change local sediment composition. Additionally, sediment composition can also be affected by other factors such as anthropic activities and (plant or animal) ecosystem engineers (Eriksson et al. 2010, van Katwijk et al. 2010, Donadi et al. 2015b).

The flooding regime and the sediment composition of the seafloor are key drivers of intertidal macrozoobenthos (Van Colen 2018) –invertebrates larger than 1 mm that live in close association with the substrate. Areas more frequently inundated will experience different environmental conditions (Beukema 1976), different predation pressure (e.g. Sanchez-Salazar et al. 1987) and may have different access to food (e.g. filter feeders). Sediment size composition determines the amount of organic matter in the seafloor (e.g. Yang et al. 2021), which influences the stability of the seafloor (Bartzke et al. 2013; but see Defew et al. 2002), and in turn the amount of food for macroinvertebrates (Christianen et al. 2017), as well as their behaviour (e.g. Wiesebron et al. 2021).

The Wadden Sea, the largest cohesive intertidal area in the world and a UNESCO world heritage site, is subject to gas extraction, which has resulted in land subsidence (Fokker et al. 2018). Macroinvertebrates of the Wadden Sea are an important part of the food web as a feeding resource for birds (e.g. Zwarts & Wanink 1993, Beukema & Dekker 2019) and fishes (Poiesz et al. 2020). Because sediment composition and flooding regime, two of the key factors that determine the occurrence and abundance of macrozoobenthic species, could be affected by land subsidence, cascading changes to the macrozoobenthic community are likely.

Gas extraction in the Dutch Wadden Sea is managed following the “hand on the tap” principle (NAM B.V 2007), meaning that if an effect on the ecosystem is detected, the gas extraction should be halted. Because of this, subsidence and its consequences are monitored through a joint effort of governmental agencies (Rijkswaterstaat), research institutions (NIOZ) and private parties (the Dutch Petroleum Company, NAM, for its acronym in Dutch). Deep subsidence occurs at a rate of ∼2 mm y^-1^ (NAM B.V 2021). However, changes in average seafloor elevation have not been detected yet (van der Vegt 2022) as subsidence has been so far compensated by increased sedimentation (Wang et al. 2018). So far, studies about the impact of gas extraction on mudflats have mostly focused on the risk of intertidal area loss (e.g. Thienen-Visser et al. 2015, Wang et al. 2018, Fokker et al. 2018, van der Vegt 2022) but changes in sediment and community composition have received less attention (but see Beukema 2002).

Detecting and measuring large-scale human impacts on ecosystems, such as those of land subsidence, is one of the big challenges in applied ecology. Ideally, such effects are examined using a before-after control-impact assessment (BACI, Green 1979). The approach has been successfully employed in many impact assessment studies (e.g. Sansom et al. 2016, Smokorowski & Randall 2017), however, there are shortcomings (e.g. Underwood 1991, Smokorowski & Randall 2017). First, the difficulty in selecting and finding appropriate and sufficient locations of control sites (Schleicher et al. 2020). Second, and related to the first problem, the observed variability between sites may be the result of spatially varying environmental variables, which confounds our ability to attribute observed changes to the anthropogenic impacts of interest (Hewitt et al. 2001). Last, the need for an impact assessment study may only be acknowledged when human activities are already ongoing, and therefore it is impossible to register the unaffected “before” conditions. Large-scale monitoring systems, when properly designed (Field et al. 2007), could overcome some of these shortcomings: sampling locations can be used for comparison if environmental variance can be controlled, and in this way the problem of confounding can be heavily reduced (discussed by Underwood 1993). This means we can detect differences between impact and control sites while accounting for differences in environmental conditions between sites.

The purpose of this study was to investigate potential effects of land subsidence caused by gas extraction on mudflat’s sediment composition and macrozoobenthic communities. To achieve this, we took advantage of thirteen years (2008-2020) of spatially comprehensive monitoring data of sediment composition and benthic intertidal macroinvertebrates. The spatial and temporal extent of these data allowed us to use it as an alternative to a BACI design. The rationale is that using the whole Dutch Wadden Sea as a reference area, and quantifying the effects of physical and biological conditions, we can control for the main processes driving sediment composition and macroinvertebrates biomass. In this way, we can detect differences between ‘within’ and ‘outside’ the subsidence area “over and above” (i.e. while controlling for) these environmental variables. Because land subsidence will change the local sedimentation regime, first, we asked whether sediment composition (median grain size and mud fraction) was different within compared to outside the area affected by land subsidence due to gas extraction. Second, as depth and sediment composition are two of the main drivers of the benthic community composition, we then asked whether the abundance of macrozoobenthos was also different within compared to outside the area affected by land subsidence. As total biomass encompasses many different species that favour different abiotic conditions, biomass might increase or decrease in the subsidence area. To clarify how the abundance of particular species might have changed, we analysed if the ratio of different species’ biomass within and outside the subsidence area changed in time. We related this ratio to the species’ favoured abiotic condition to identify species that may be benefitted or hindered by land subsidence. Additionally, because the area affected by subsidence is much smaller than the rest of the Wadden Sea, we replicated these analysis using areas unaffected by land subsidence to better understand how the natural variability of the Wadden Sea could affect our results.

## Materials and Methods

### Study area

The Wadden Sea extends for 8000 km^2^ from the north of the Netherlands to the east of Denmark (53° N Lat, Fig 1). It has a bidaily mesotidal regime with a range of 1.4 to 3.4 m across the system (Postma 1982). The soil is composed by sand and mud (Oost 1995). Its macrozoobenthic community is dominated by bivalves and polychaetes, with *Cerastoderma edule, Mya arenaria* and *Arenicola marina* being the most abundant species it terms of biomass (Compton et al. 2013). Gas extraction started in the area of Ameland in 1986 (Fokker et al. 2018). Since then, measurements of land subsidence have been taken by the NAM through various methods: leveling (comparing elevation against the national height reference network, Fokker et al. 2018), every 4 years since 1986, surveying with GPS, once a year since 2006, and interferometric synthetic aperture radar measurements (Insar, on land only) once a year since 1997 (NAM B.V 2021). These measures were modeled by NAM to determine the contours and depth of deep land subsidence. For this study we use the contours of 1 cm of deep subsidence registered since 2006 (Fig. 1) (NAM B.V 2021).

**Fig 1:**
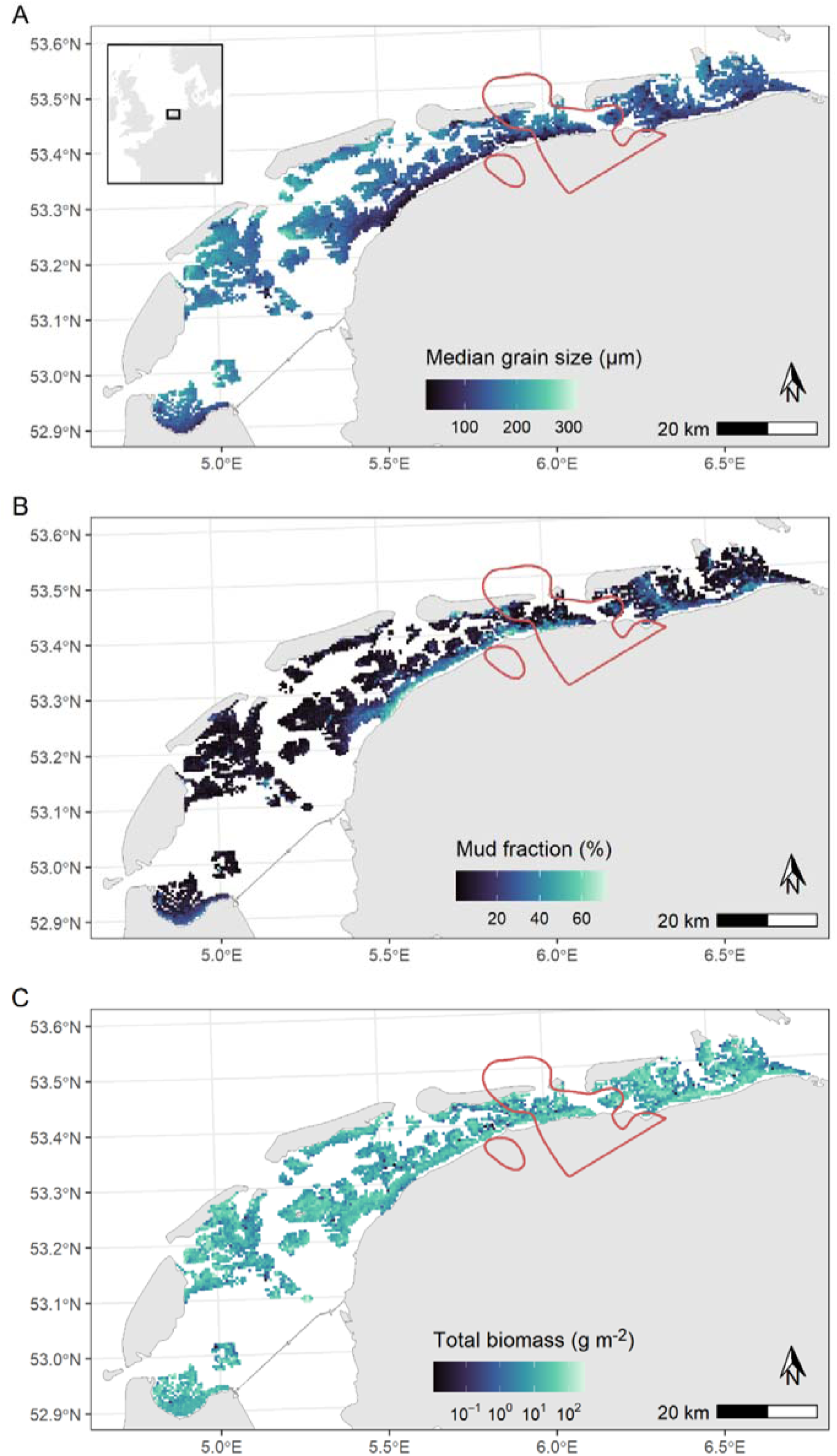
Synoptic intertidal benthic survey (SIBES) sampling stations (each point is a station) and subsidence contour in red (1 cm of subsidence between 2006 and 2020). **A**) Average (2008-2021) median grain size, **B**) average mud fraction, and **C**) average total macroinvertebrate biomass (measured as ash-free dry mass) at each sampling station.

### Survey

Between 2008 and 2020 the macrozoobenthic community and sediment composition of the Wadden Sea were sampled for the synoptic intertidal benthic survey (SIBES). Every year, between June and October, we aimed to visit 4511 stations (Fig. 1) of which 78% were located on a regular (500 m) grid design and the rest were randomly distributed along the grid (Bijleveld et al. 2012). Due to logistic constraints, each year between 1441 and 4209 stations were effectively visited (see distribution of samples per station in ESM 1). 528 stations were within the contour of subsidence from which we obtained 3866 samples during the entire studied period. Stations are visited either by foot or using a small rubber boat, depending on tidal and sediment conditions.

Sediment samples were taken with a 33 mm diameter core to a depth of 4 cm and conserved at –20° C. In the laboratory, samples were freeze-dried for 96 h, homogenized with mortar and pestle, weighted and put in 13 ml auto-sampler tubes with degassed reverse osmosis water. Samples were not treated to remove calcium or organic material. Sediment size composition was characterized using a particle size analyzer (Coulter LS 13 320) that measures grain sizes between 0.04 and 2000 µm in 126 size classes through laser diffraction and polarization intensity differential scattering technology. For each sample, the median grain size and mud fraction (volume percentage of particles smaller than 63 µm) were obtained.

For the benthic community, a core sample of 0.017 m^2^ was taken to a depth of ∼25 cm and sieved through a 1 mm round mesh. All animals retained were fixed in formalin (4% solution) except for bivalves larger than ∼1 cm that were frozen. In the lab, individuals were identified to the finest taxonomic level possible, and ash free dry mass (afdm) was measured to 0.1 mg. For a more complete description of the methods see Compton et al. (2013).

### Covariates

We identified 10 environmental variables known to be of importance for the distribution of sediment and/or macroinvertebrates (Table 1). Elevation and current shear stress determine the amount of energy in a particular place and therefore are associated with sediment composition. Elevation is also an important driver of macroinvertebrates distribution (e.g. Kraan et al. 2010, de Jong et al. 2015). Strong shear stress and waves may dislodge benthic invertebrates, so we also expect these variables to affect their total biomass (e.g. Anta et al. 2013, Malakauskas et al. 2013, but see Donadi et al. 2015a). We used elevation data from bathymetric maps of the Dutch Wadden Sea that are updated every five years and made available by Rijkswaterstaat. The distribution of mean bottom shear stress due to currents in the Dutch Wadden Sea was obtained from Folmer et al. (2016) who calculated it using the General Estuarine Transport Model (Burchard & Bolding 2002).

**Table 1:**
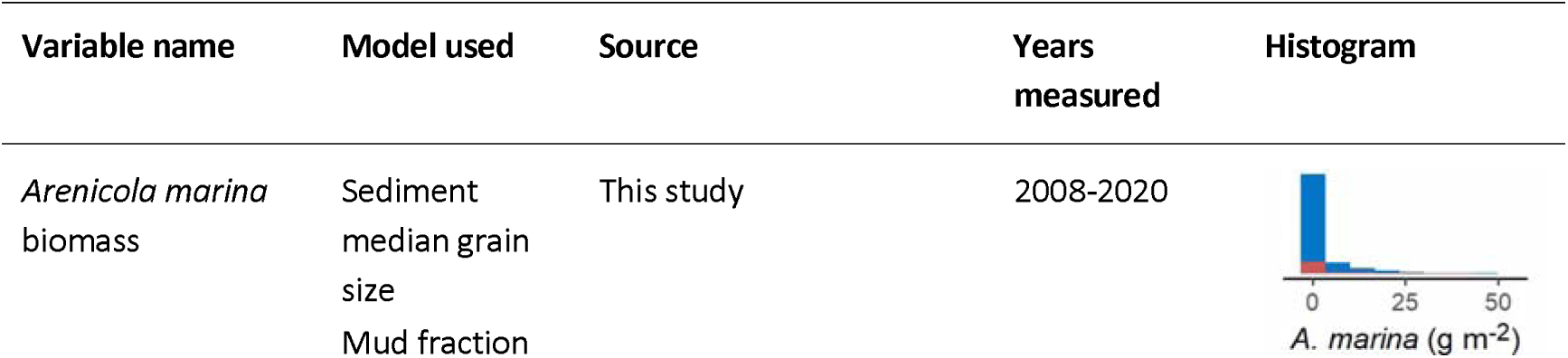
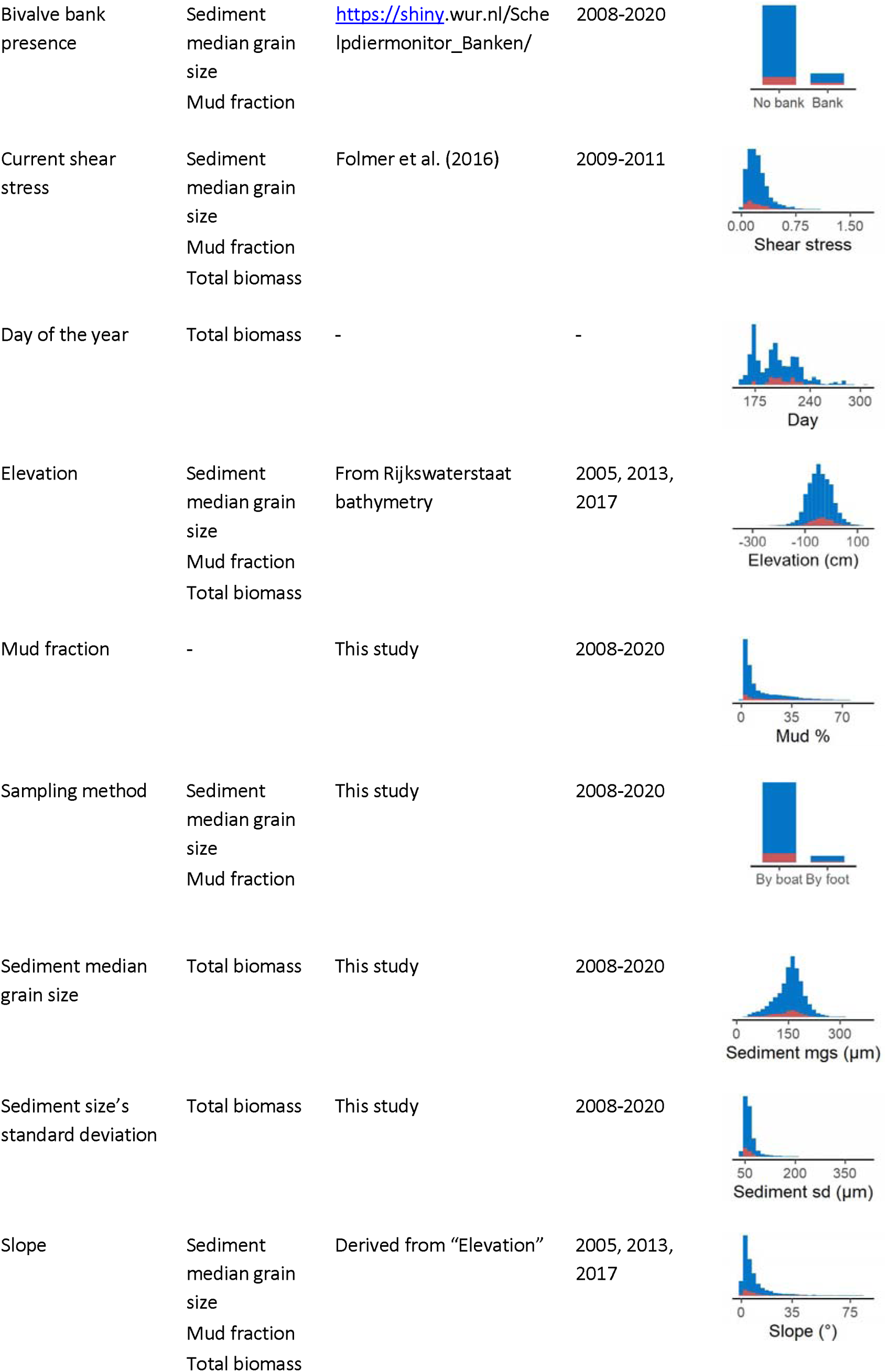
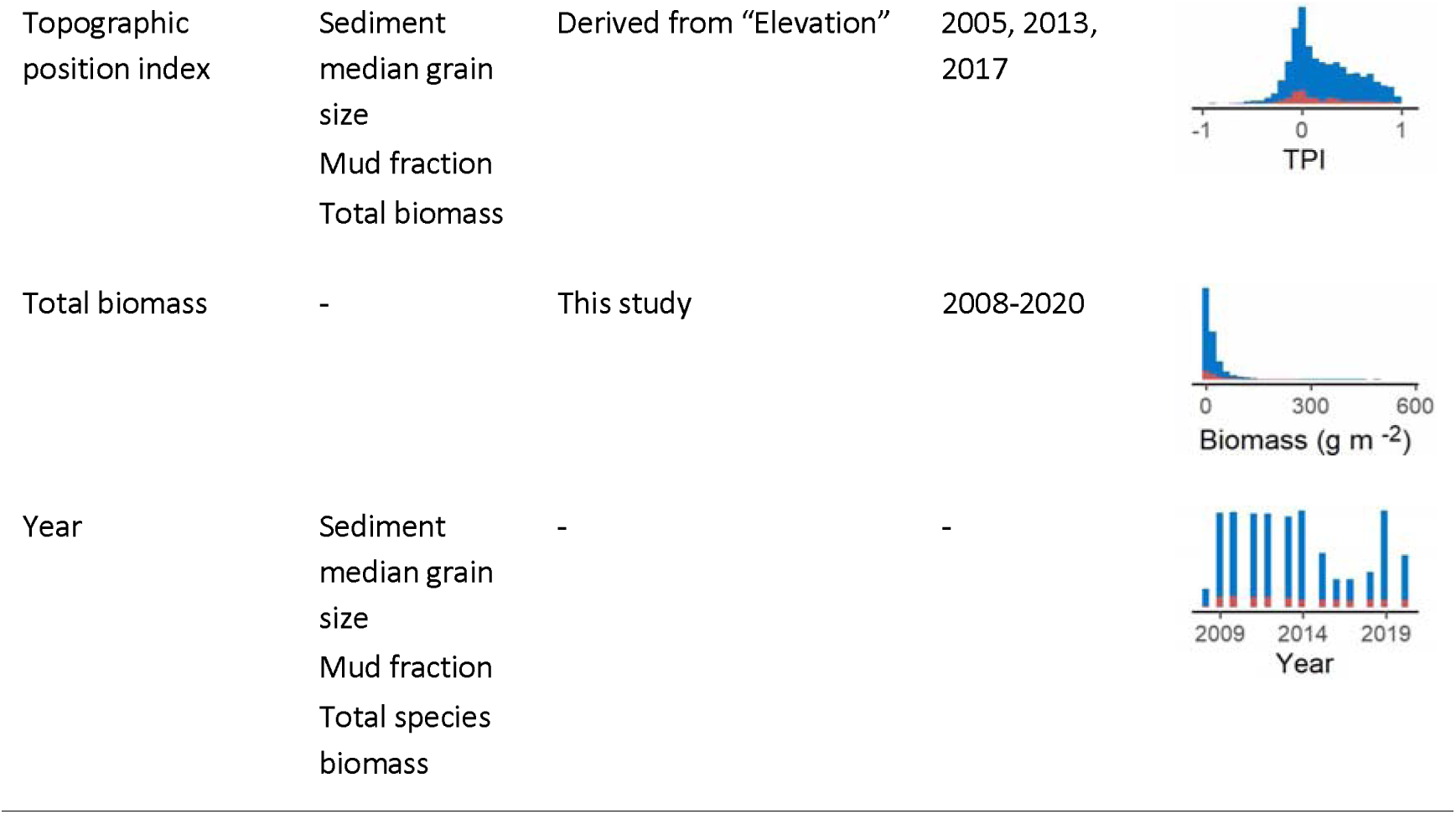
Alphabetically ordered variables that were included in the models fitted to sediment median grain size, mud fraction, and total biomass data. Source of the data, year of measurements, frequency distributions of the data and in which model these variables were included is shown. The colors in the frequency plots indicate outside (blue) and inside (red) the subsidence area.

In soft sediment systems, seafloor topographic features are correlated with different sediment composition (e.g. Trentesaux et al. 1994, Mestdagh et al. 2020). Different benthic communities are also associated with different topographic features (e.g. Jansen et al. 2018, Mestdagh et al. 2020). For this reason, we included slope, topographic position index and topographic ruggedness as covariates to the models for median grain size, mud fraction and biomass. We derived this information from the bathymetry. Slope was calculated using function ‘slope’ from R package starsExtra (Dorman 2021), topographic position index using function ‘wbt_relative_topographic_position’ from R package whitebox (Qiusheng 2019) and a kernel of 79 m. Ruggedness was calculated through function ‘wbt_ruggedness_index’ also from R package whitebox (Qiusheng 2019).

Sediment composition strongly influences the benthic community (e.g. Kraan et al. 2010, de Jong et al. 2015), therefore, median grain size, mud fraction and the standard deviation of sediment size were included in the biomass model. Day of the year was also included in the biomass model to account for periods dominated by growth or mortality.

The lugworm, *Arenicola marina*, the blue mussel *Mytilus edulis* and the Pacific oyster *Crassostrea gigas* are important ecosystem engineers in the Wadden Sea (Volkenborn et al. 2007, Donadi et al. 2015b). *A. marina* removes the sediment resuspending the smaller particles. Oysters and mussels, when filter-feeding, catch the smaller particles suspended in the water column and fix them in the sediment when excreting. The biomass of *A. marina* and the presence of a *M. edulis-C. gigas* bank were used as covariates in the median grain size and mud fraction models. However, because the effect of *A. marina* over the sediment is small (∼13%) and only surficial (Volkenborn et al. 2007), we also ran additional versions of the models not including this covariate (ESM 2). *A. marina* biomass was determined through SIBES samples as described in the ‘survey’ section. Mussel and oyster banks in the Wadden Sea are mapped every year (Troost et al. 2022). Because the effects of such bivalve banks on sediment extends beyond the bank itself (Kröncke 1996), we added a 300 m buffer around the bivalve beds contours and classified samples within these contours as “within bivalve bank”.

Finally, the sampling method (by foot or by rubber boat) was also considered a potential source of variation for median grain size and mud fraction. Samples are taken by boat when the mudflat is inundated and they are taken by foot only when the mudflat is exposed. This difference between inundation conditions might affect the composition of top layer of the sediment, so we controlled for this by considering the sampling method as a covariate.

### Spatial modelling

We modeled three response variables: sediment median grain size (μm), mud fraction (%) and total macroinvertebrate biomass (afdm, g m^-2^), each using an appropriate error distribution. We used a Gaussian distribution for median grain size and a beta distribution with a logit link function for mud fraction (Douma & Weedon 2019). As all the samples contained detectable biomass, a Gamma distribution with a log link was used for total biomass (Zuur et al. 2019).

Covariates described in the previous section and Table 1 were included in the models as fixed-effects. To avoid numerical estimation problems, all covariates were standardized by subtracting their mean and dividing by their standard deviation. Collinearity between variables was assessed by computing variance inflation factors (VIF) with function ‘vif’ of car R package (Fox et al. 2011). We sequentially dropped the covariates with the highest VIF until all presented VIF < 2 (Zuur et al. 2010). In this way, topographic ruggedness was dropped from the models. We tested if the relationship between the link function and each covariate was linear by running generalized linear mixed models and visually inspecting the plots of Pearson’s residuals against each predictor variable. If the residuals showed a residual trend along the predictor variable, this indicated that the relationship between the link function and the predictor was non-linear, and thus, we used a smoother for said covariate in the final model (Zuur et al. 2019).

Subsidence was included in the model as a dichotomous variable: each observation was classified as within or outside the subsidence area. We did this because 38% of our observations within the subsidence area fell inside the 1 cm contour, and therefore we did not have enough information to analyze its effect as continuous. We assessed whether the effect of subsidence changed in time by adding an interaction term with ‘year’. Priors for all fixed effects were set to a mean of zero and a precision of 1. This high prior weight around zero results in the need of more data support for the posterior of a parameter to be different from zero, which makes our results more conservative.

To account for spatial autocorrelation between observations that cannot be explained by the available covariates, a stochastic partial differential equation model (SPDE, Lindgren et al. 2011) was used. In this approach, spatial pseudoreplication is dealt with by adding a spatially correlated random intercept that follows the distribution of a Gaussian field with mean zero and the correlation between two locations is given by a Matérn function (Zuur et al. 2019). To estimate the random intercepts at different points, a mesh of triangles is constructed for the study area and used to estimate the spatially and temporally structured random component (SPDE specifications in ESM 3). To prevent the SPDE from absorbing too much of the variability of the model, penalized complexity priors were added. These priors were set following Fuglstad et al. (2019) and Zuur et al. (2019). To account for temporal autocorrelation from different survey years, we assessed three temporal correlation structures for the spatial random field: no correlation, “AR1” and “replicated” (Zuur et al. 2019). We ran the full models using each of the temporal correlation structures and kept the one with lower DIC (see ESM 3).

The models were validated by inspecting their Pearson’s residuals and checking for homogeneity of variance along the fitted values, and no residual patterns in the residuals when plotted against covariates (Zuur et al. 2019). Residual spatial autocorrelation was assessed by plotting residuals along x and y coordinates (Zuur et al. 2019).

Our main objective with this study was to test if there was a difference in the response variables depending on subsidence, therefore we focused on hypothesis testing rather than on model selection (Fieberg 2022). In Bayesian statistics the uncertainty around any parameter can be estimated (Fieberg 2022) so we took advantage of this trait to estimate 95% confidence intervals around the parameter of each covariate, including subsidence and its interaction with time. If zero lies within the confidence interval of a parameter, it means we cannot distinguish the effect of that variable from zero with 95% confidence. Therefore, we considered that subsidence had an effect on sediment or biomass if zero was outside the range of the confidence intervals for the subsidence or interaction term (Fieberg 2022).

We ran additional analyses to assess if differences similar to those found in the original models arouse by chance. In these analysis we compared a group of samples not affected by subsidence to the rest of the Wadden Sea. We called these groups of samples “fake subsidence areas”. Because each “fake subsidence area” should be unique to avoid pseudoreplication, the number of potential areas was limited. We selected five “fake subsidence areas”, each centered in a different tidal basin of the Wadden Sea and with similar sample sizes as the real subsidence area (see details in ESM 4). We ran models for median grain size, mud fraction and total biomass with the same specifications as the original ones, but using the “fake subsidence” area instead of the real subsidence area, and using the rest of the Wadden Sea except for the real subsidence area as the “no subsidence area”. If the effect of “fake subsidence” and the interaction between “fake subsidence” and year is not different from zero, this would support the differences originally found.

### Species-specific effects of land subsidence

Total biomass is an aggregate of different species which might be differently affected by subsidence due to differences in natural history and ecology. To analyze if particular species are driving the observed biomass patterns, we identified the more abundant species in terms of biomass within the subsidence area, and plotted their contribution to total biomass within and outside the subsidence area. These species are *Arenicola marina, Cerastoderma edule, Lanice conchilega, Mya arenaria* and *Mytilus edulis* and together they represent more than 90% of total biomass within the subsidence area each year.

To understand how species with different habitat use are affected by subsidence, we compared population trends within and outside the subsidence area for a larger group of species (species selection methods in ESM 5). To classify species’ habitat use, for each species, we extracted the median of the ‘median grain size’ and the median of the elevation range in which they occurred. Hereafter we call these median values ‘sediment use’ and ‘elevation use’.

To detect differences between population trends, we calculated the ratio between the average biomass within and outside the subsidence area for each species and year. We used a linear mixed model with log(‘biomass ratio’) as response variable, ‘elevation use’ and ‘year’ as continuous fixed effects, and ‘species’ as random slope and random intercept. We did not include ‘sediment use’ in the model to avoid collinearity with ‘elevation use’ (Pearson’s correlation: –0.72, *P* < 0.001). We applied a likelihood ratio test to select the best random structure and AIC together with 95% parameter confidence interval to select the best model relative to fixed effects (Zuur 2009). As an additional control, we ran the same analysis, but using the “fake subsidence areas” instead of the subsidence area, in the same way as described for the previous section (see details in ESM 4). If the log biomass ratio between the “fake subsidence” area and the rest of the Wadden Sea remains constant between years, this would support the validity of the differences originally found.

## Results

### Median grain size

The average median grain size of all samples was 154 ± 41 µm (mean ± sd, Fig 1A). The mixed-effects model fitted to median grain size revealed a positive effect of current shear stress and topographic position index, and a negative effect of the presence of bivalve banks (Fig 2A). The local elevation and biomass of *Arenicola marina* had non-linear effects on median grain size (Fig 2A, ESM 6). The model revealed a significant effect of the interaction between the subsidence term and the year term on median grain size (Fig 2A). This meant that while there was no evidence for a positive or negative trend in sediment median grain throughout the Wadden Sea, within the subsidence area, median grain size has been decreasing (Fig 2B). Taking *A. marina* out of the model did not qualitatively change the results (ESM 2). At the end of the study period, the model estimated that median grain size within the subsidence area was on average 16 μm smaller than outside (Fig 2B). The “fake subsidence” analysis showed that comparing areas of similar size to the rest of the Wadden Sea, we do not find differences in median grain size (ESM 4).

**Fig 2:**
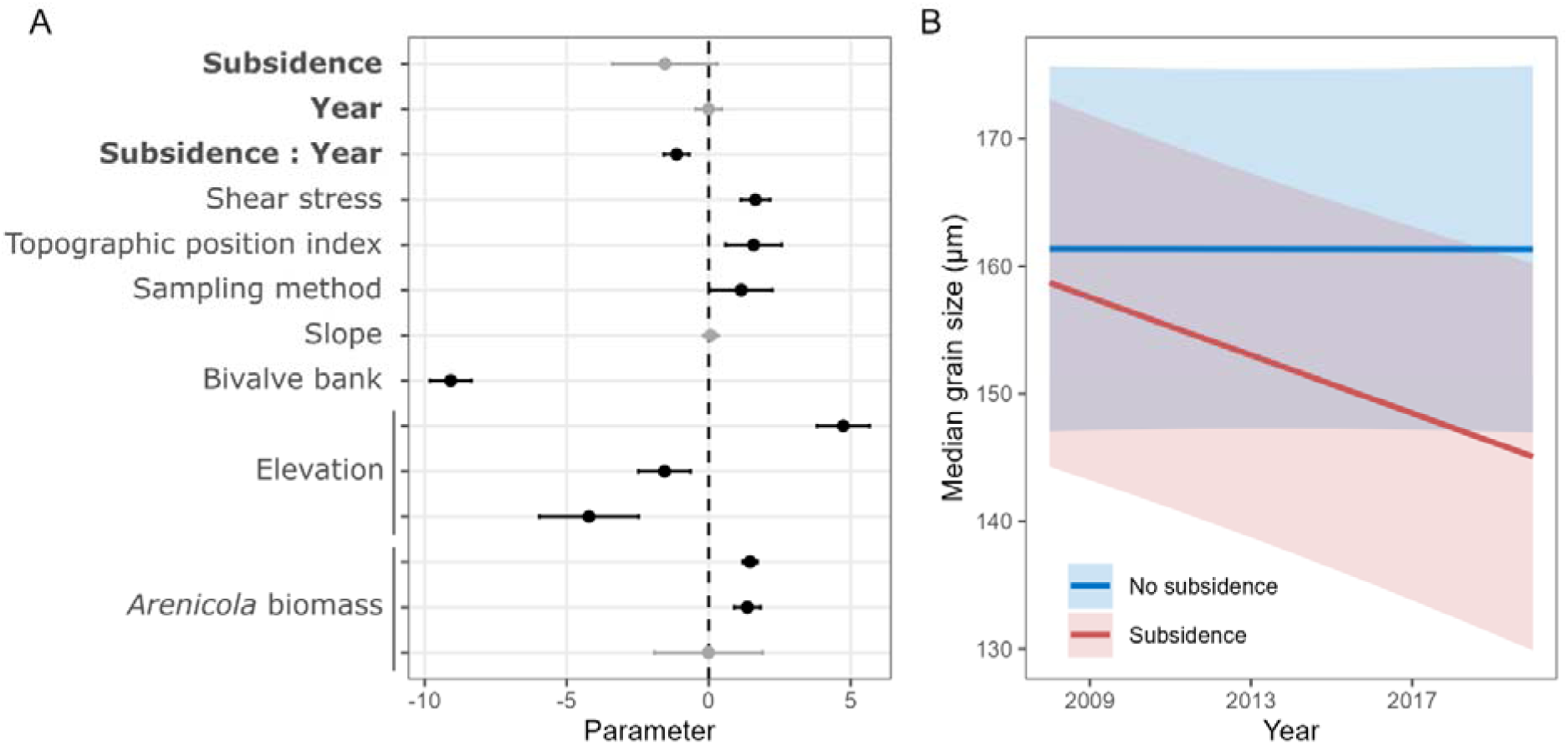
Results for the sediment median grain size model. **A**. Parameter estimates for covariates used in the model (mean ± 95% CI). Covariates for which a smoother was applied have more than one parameter. Points and lines in black show effects that are different from zero with 95% confidence. **B**. Predictions for median grain size (mean and 95% CI), within and outside the subsidence area throughout the study period.

### Mud fraction

On average, samples contained 0.13 ± 0.13 fraction of mud (mean ± sd, Fig 1B). Mud fraction was negatively affected by most of the covariates assessed, but positively affected by bivalve bank presence (Fig 3A). We found a positive effect of subsidence on the mud fraction (Fig 3A) that remained constant in time (Fig 3B). Taking *A. marina* out of the model did not qualitatively change our results (ESM 2). On average, the mud fraction within the subsidence area was 0.03 (3%) higher than in the rest of the Wadden Sea. The “fake subsidence” analysis showed that comparing areas of similar size to the rest of the Wadden Sea, we do not find differences in mud fraction comparable to those found between the subsidence area and the rest of the Wadden Sea (ESM 4).

**Fig 3:**
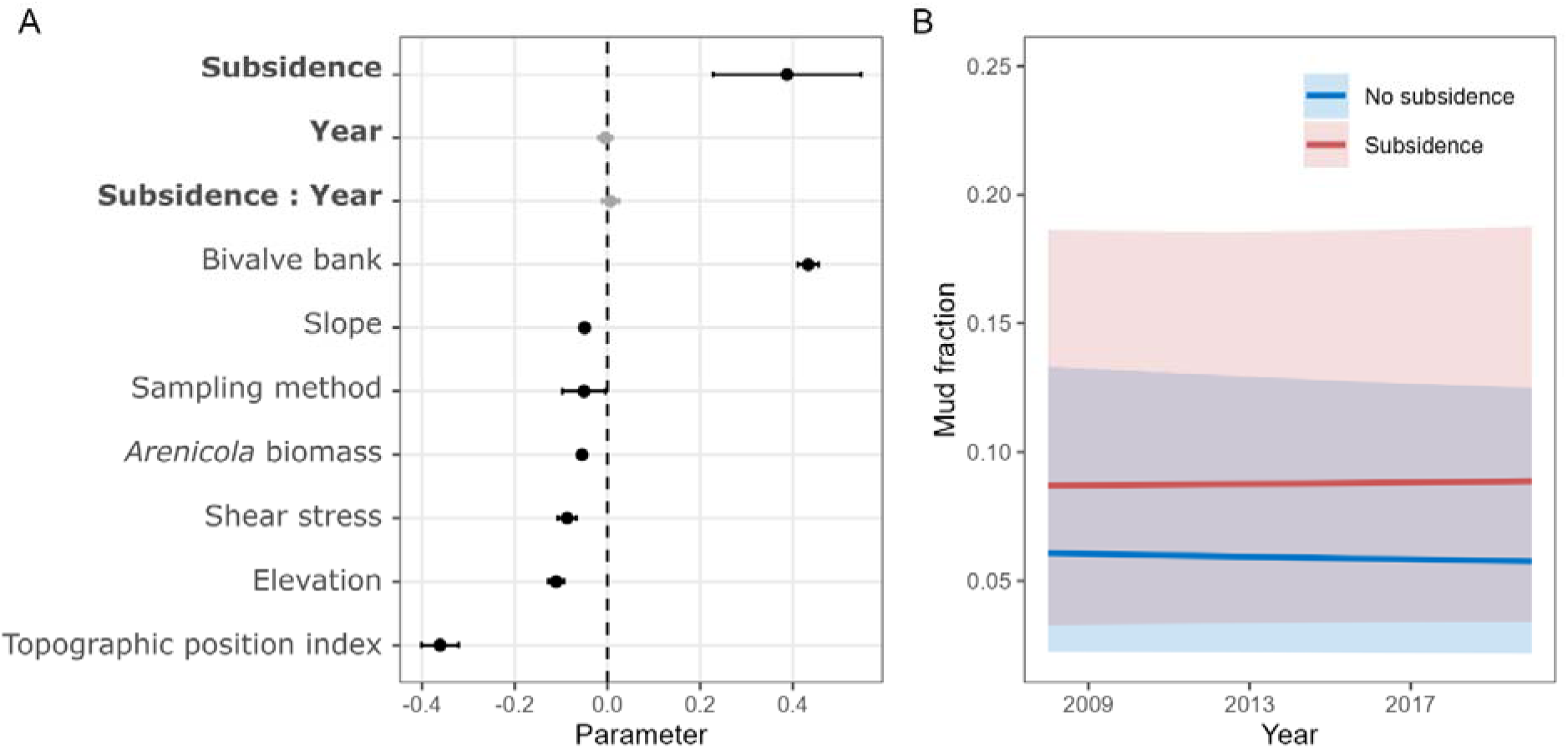
Results for the sediment mud fraction model. **A**. Parameter estimates for covariates used in the model (mean ± 95% CI). Points and lines in black show effects that are different from zero with 95% confidence. **B**. Predictions for mud fraction (mean and 95% CI), within and outside the subsidence area throughout the study period.

### Biomass

We found that total biomass was on average 24.5 ± 45.3 g m^-2^ (mean ± sd, Fig 1C). Biomass was negatively affected by topographic position index current shear stress and slope (Fig 4A). Elevation, day of the year, median gran size and the standard deviation of sediment grain size had non-linear effects on biomass (Fig 4A, ESM 6). We did not find an effect of subsidence on total biomass of the macroinvertebrate community (Fig 4A,B). The “fake subsidence” analysis showed differences in the total biomass of two areas compared to the rest of the Wadden Sea (ESM 4).

**Fig 4:**
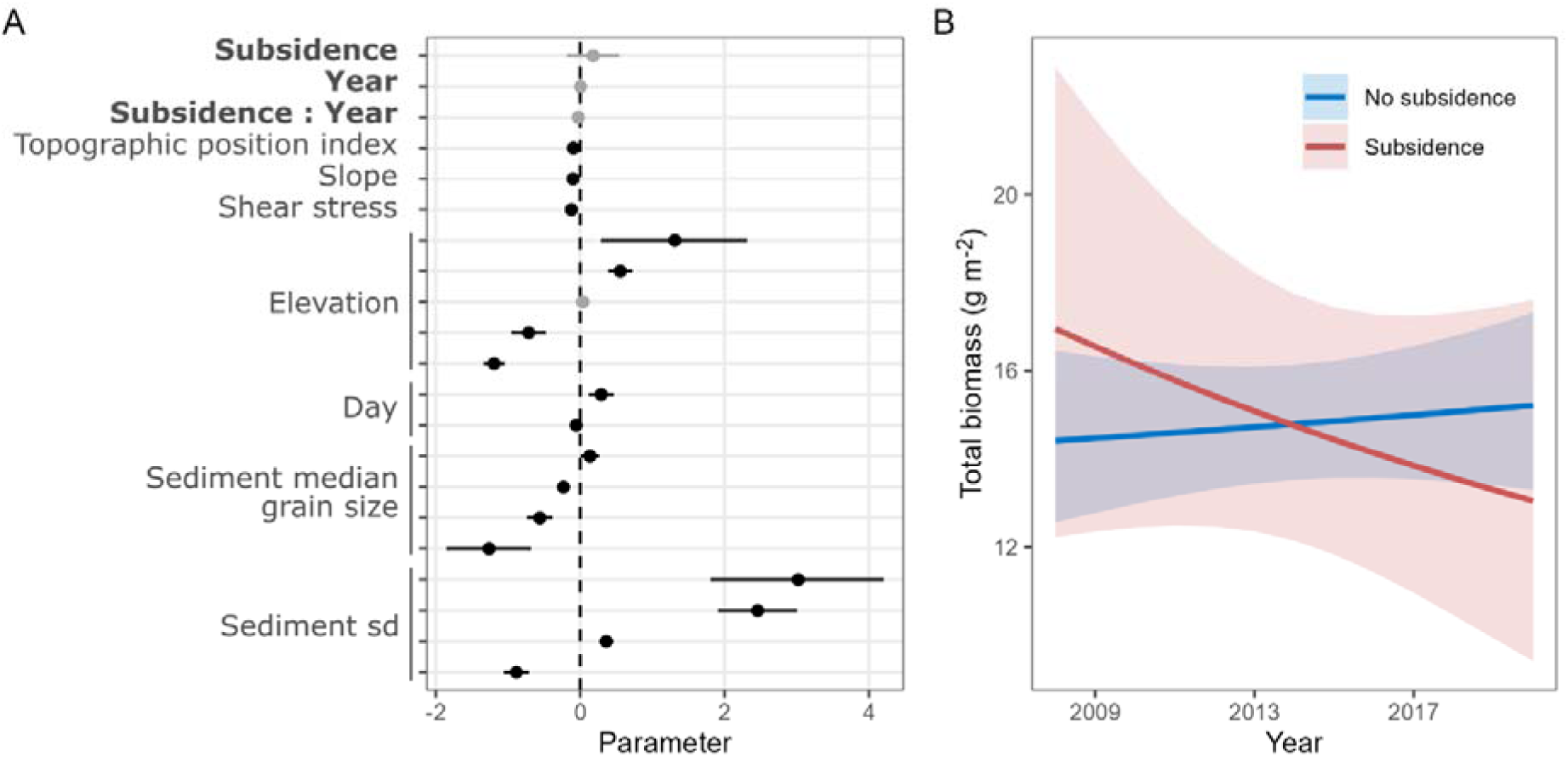
Results for the total biomass model. **A**. Parameter estimates for covariates used in the model (mean ± 95% CI). Covariates for which a smoother was applied have more than one parameter. Points and lines in black show effects that are different from zero with 95% confidence. **B**. Predictions for biomass (mean and 95% CI), within and outside the subsidence area throughout the study period.

### Species-specific effects of land subsidence

The most abundant species (in terms of biomass) within the subsidence area were *Arenicola marina*, *Cerstoderma edule*, *Lanice conchilega*, *Mya arenaria* and *Mytilus edulis*. These species also represented an important fraction of biomass outside the subsidence area (Fig 5), with other species only contributing up to 5 ± 1% of total biomass. From 2008 to 2019, *C. edule* was the species with highest average biomass in both areas. *C. edule* represented the 56% of total biomass within the subsidence area in 2008, but only the 40% outside. In 2020, *C. edule* was surpassed by *A. marina* in both areas, and by M. arenaria outside the subsidence area (Fig 5).

**Fig 5:**
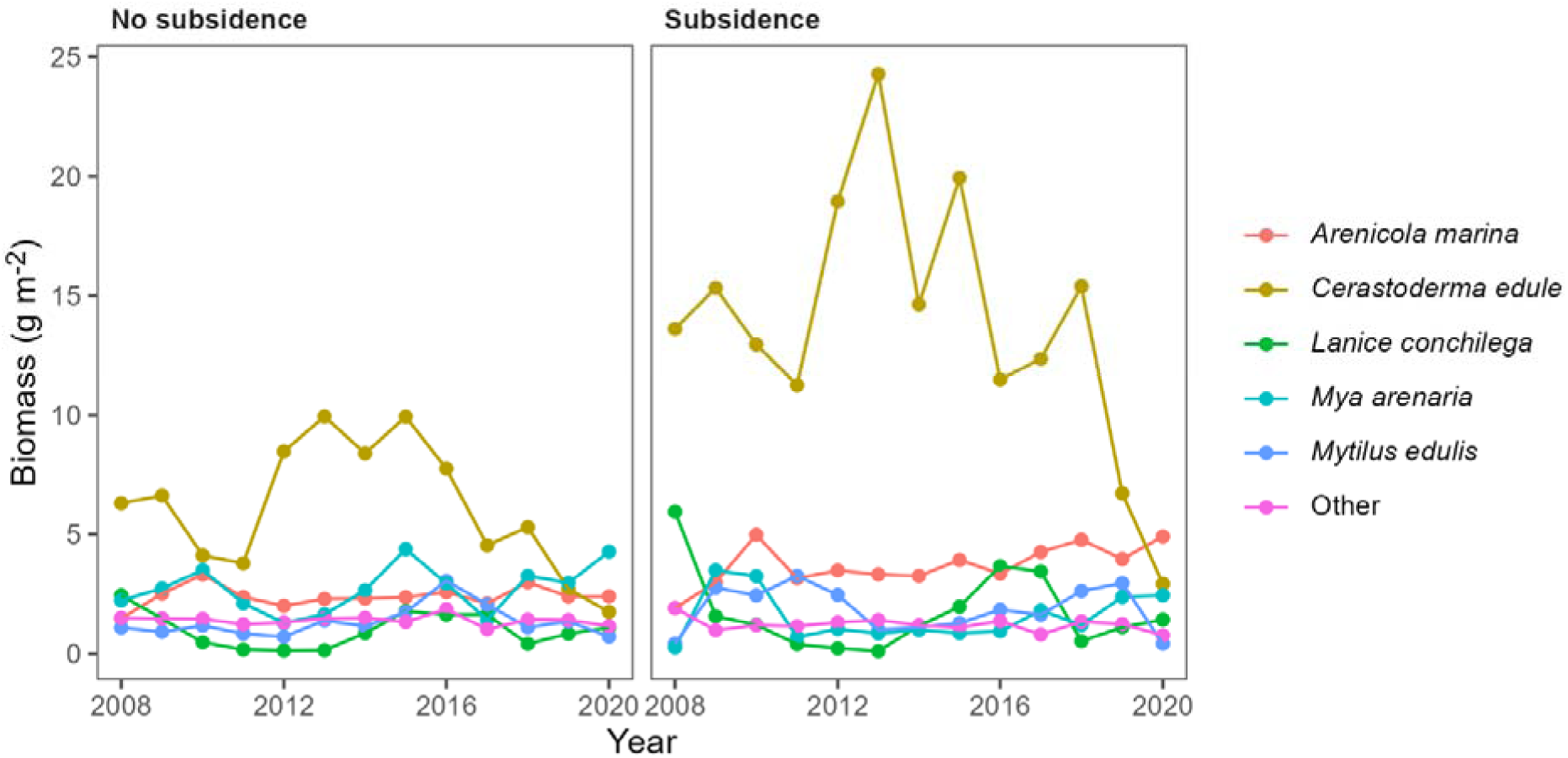
Species biomass within and outside the subsidence area. The portrayed species are the ones with higher contribution to biomass within the subsidence area.

With the mixed-effects model, we found differences in population trends within and outside the subsidence area, and these differences depended on the ‘elevation use’ of species (Fig 6). For species that use lower areas, populations have increased in biomass within relative to outside the subsidence area. On the contrary, for species that use higher areas, populations have been slightly decreasing within relative to outside the subsidence area (Fig 6). However, when running the “fake subsidence” analysis, we also found differences in population trends when comparing “fake subsidence” areas with the rest of the Wadden Sea (ESM 4). This means that the differences in species composition found between the subsidence area and the rest of the Wadden Sea might be are a reflection of the natural variability of the ecosystem.

**Fig 6:**
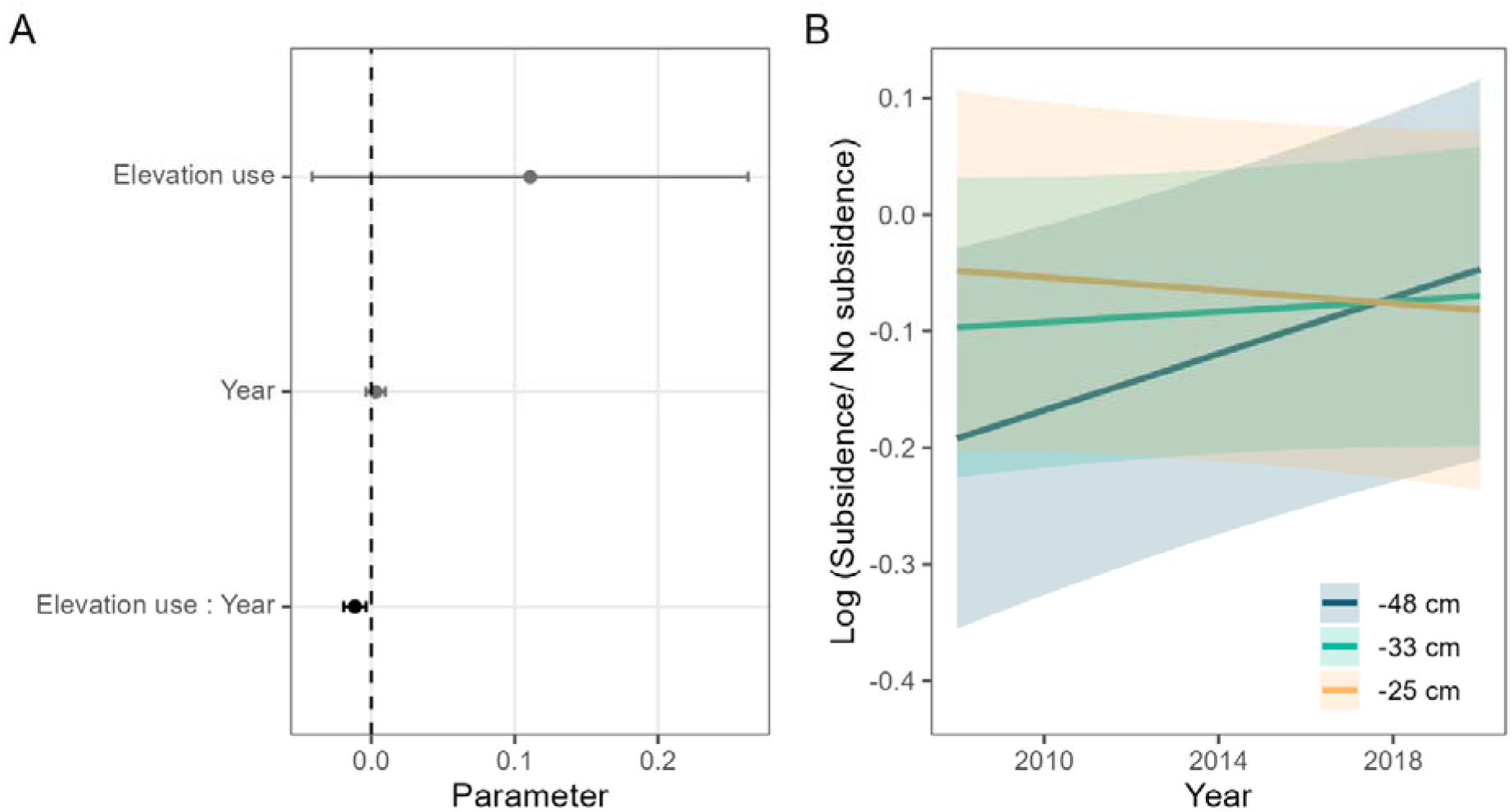
Results for the population trends model. **A**. Parameter estimates for covariates (mean ± 95% CI). Points and lines in black show effects that are different from zero with 95% confidence. **B**. Mean and 95% confidence intervals of the ratio of biomass within and outside the subsidence area throughout the study period.

## Discussion

We found differences in sediment composition within the land subsidence area relative to outside. After accounting for confounding environmental variables, median grain size within the subsidence area decreased by 10% during the study period and the mud fraction was 3 percentage points higher throughout.

These changes do not seem to significantly affect the total biomass of the macroinvertebrates living there. We detected a change in species composition with an increase in the biomass of species that use deeper habitat and a decrease of species that use a shallower habitat. However, we could not discard these changes being part of the greater variability of the Wadden Sea.

Subsidence potentially occurs in an area with unique characteristics within the Wadden Sea, and any change in the subsidence area could be caused by other natural or anthropogenic processes there. For example, the land subsidence area is close to the Lauwerszee, that has been closed-off in 1969, changing the sediment transport dynamics of the system (Elias et al. 2012), and one of the inlets within the subsidence area moves in a cyclic way, changing and recovering its shape every 20-40 years (Oost 1995, Elias et al. 2012). The potential uniqueness of the area raises the issue of a lack of a proper control in this study, a problem inherent to nearly all large-scale ecological field studies. To attribute causation, a BACI design is necessary (Green 1979), however, subsidence has been going on since the eighties, long before we started collecting the data. For this reason, using the initial conditions (i.e. data from 2008) as the control would be incorrect. Median grain size, mud fraction and biomass are, nonetheless, not randomly distributed in space, rather their distribution results from environmental and biological processes that we aimed to capture in the statistical model with the covariates (like elevation and current shear stress). Because the range of all the covariates within the subsidence area fall within the range of the whole Wadden Sea’s (see histograms in Table 1), we can thus attempt to detect differences “over and above” environmental differences between the subsidence area and the rest of the Wadden Sea assuming that the underlying processes (mechanisms) are similar. On top of this, the addition of a spatial latent field with a temporal structure allows us to at least partially control for spatially and temporally structured errors caused by historic events (e.g. the closure of the Lauwerszee) and missing covariates (e.g. suspended sediment loads). Of course, we cannot discard alternative explanations to the differences we observe, but even after accounting for confounding environmental variables and residual spatiotemporal patterns, we still observed differences in median grain size and mud fraction within and outside the subsidence area. Supporting these results, we could not detect differences between any other particular area and the rest of the Wadden Sea, showing that the observed differences are likely not due to the natural variability of the Wadden Sea.

Studies on effects of land subsidence in coastal systems have so far mainly focused on the direct effects of surface subsidence: loss of elevation and consequent land loss (e.g. Kolker et al. 2011, Al Mukaimi et al. 2018), and its relationship with the sediment accreting capacities of the environment (Reed 2002, Al Mukaimi et al. 2018). Most of them are carried out on salt marsh ecosystems. We have not found any other study about the composition of the sediment deposited in intertidal systems after land subsidence (but see Molinaroli et al. 2009, Al Mukaimi et al. 2018). In terrestrial systems, the effects of land subsidence on soil composition have been explored generally by comparing soil composition at different areas of the subsidence bowl (top, slope, bottom) with control areas (Tripathi et al. 2009, Qing-jun et al. 2009, Guo et al. 2018, Ma et al. 2019, Vishwakarma et al. 2020). The lack of measurements from before an area is impacted is extensive for this kind of studies. We found only one study that characterized the soil composition before the subsidence occurred (Dejun et al. 2016) and one studying landscape scale effects through maps made before the impact (e.g. Quanyuan et al. 2009). This speaks about how difficult it still is to achieve a proper experimental design to assess large scale anthropic impacts on the environment.

### Subsidence and sediment composition

Subsidence has not affected the average bathymetry and morphology of the tidal flats (van der Vegt 2022), which implies that sedimentation was so far able to keep up with subsidence. Sand entering the Wadden Sea mainly comes from the erosion of the ebb-tidal deltas which are in turn supplied by coastal nourishments (Elias et al. 2012). Mud is transported along the Dutch Coastal zone by the residual north-easterly current (van der Hout et al. 2015) and enters the Wadden Sea through the tidal inlets. Within the subsidence area, median grain size was decreasing, and the overall mud fraction was smaller throughout the study period. Modelled deep subsidence is bowl-shaped (Fokker et al. 2018), with deeper parts in the centre and shallower parts on the edges of the subsidence contours. Sediment tends to be smaller in troughs (e.g. Mestdagh et al. 2020) so we hypothesize that the bowl-shaped area could therefore promote the sedimentation of particles smaller-than-expected based on the environmental characteristics, resulting in finer sediment composition in this area. Furthermore, previous studies have shown that the role of mud in the sediment infilling of the Wadden Sea has increased in the last century (Colina Alonso et al. 2021). We hypothesize that the larger sedimentation rates that subsidence promotes, together with the more prominent role of mud in infilling the Wadden Sea might result in the smaller grain sizes observed in this study. It is possible then, that we are detecting the effects of subsidence on sediment composition, while changes in bathymetry or morphology might be too subtle or compensated too fast to be detected.

Mud fraction was slightly higher within the subsidence area in the beginning of the study and remained stable during it, showing a different pattern than median grain size. This indicates that the decrease in median grain size is possibly due to an increase of small sand rather than an increase in mud fraction. We hypothesize that the different patterns observed in mud fraction and median grain size might be the result of mud and sand deposition being governed by independent processes: the settling velocities of the two types of particles differs by an order of magnitude, and while suspended sand concentration depends on local hydrological conditions, suspended mud concentration depends on supply (Colina Alonso et al. 2022). Furthermore, because subsidence has been going on since the eighties (Fokker et al. 2018), we hypothesize that these differences could be the result of a difference in speed where the effects on mud fraction have plateaued, while the sand fraction is still being affected.

### Subsidence and macrozoobenthos

We did not find changes in macrozoobenthos total biomass within the subsidence area relative to outside, but we found that species that use deeper areas are increasing within the subsidence area, relative to outside. However, we also found a significant interaction between depth use and year in one of the five “fake subsidence” controls. This means that the relative biomass of species is also changing in other areas of the Wadden Sea. Therefore, we cannot discard the hypothesis that the differences in species composition found between the subsidence area and the rest of the Wadden Sea are a reflection of the natural variability of the ecosystem. Due to the interdependency of elevation and sediment composition, species that use deeper areas are generally those that can be found in coarse sediment. With this study, we therefore cannot tell if species are choosing for a specific sediment or for a specific elevation. Changes in the elevation of the mudflats occur throughout the Wadden Sea, with certain areas getting higher and others lower as a result of natural and anthropic processes (Elias et al. 2012). This is also true for the subsidence area that has gained up to ∼7 cm per year between 2010-2021 in some areas, and lost a similar amount in others, while its average elevation remained mostly unaffected (van der Vegt 2022). We hypothesize that changes in local elevation might transform the suitability of areas for different species resulting in the increase and decrease of species depending on their depth use within the subsidence area, but also in at least one other (“fake subsidence”) area in the Wadden Sea. Further studies focusing on the changes in functional groups composition might shed light on the process behind this pattern.

### Other sources of subsidence in the Wadden Sea

There are other areas that were or are subject of anthropogenic subsidence in the Wadden Sea (Fokker et al. 2018). Gas extraction at the west of the Wadden Sea (in Zuidwal), took place between 1988 and 2020, resulting in 69.5 mm of subsidence at the deepest point up until 1997 (Rommel 2004) and less than 0.01 mm y^-1^ by 2018 (Fokker et al. 2018). Salt solution mining (near Harlingen) started in 2020 and is ongoing. It is expected to result in 106 cm of subsidence by 2052 in the deepest area (Fokker et al. 2018). There is gas extraction below Groningen since 1963, and the resulting subsidence is partially extending to the Wadden Sea (Fokker et al. 2018). Because of the difference in rates, start and finish dates, and the way in which these events were or are being monitored (Fokker et al. 2018), we do not consider the information available comparable in order to include it in this study. Additionally, subsidence might not have the same effect in the different areas. For example, while here we address subsidence in a sediment accommodation-limited basin, subsidence in Zuidwal occurs in a sediment deposition limited basin (Wang et al. 2018), which means it’s likely that it will not be filled out by sedimentation as easily, and changes in sediment composition will likely be different. The areas of the Wadden Sea affected by subsidence from Zuidwal and Groningen are much smaller than the subsidence area studied here (e.g. ∼1700 samples within Zuidwal, see Hoeksema et al. 2004) and therefore, we do not expect them to drive the patterns in average sediment compositions we observe in the rest of the Wadden Sea.

### Management implications

The Wadden Sea as a UNESCO heritage site is a protected area. The gas extraction near Ameland is managed following the “hand on the tap” principle, meaning that “if the natural values of the Wadden Sea are compromised”, gas extraction must be reduced or even ceased. Intertidal area loss is the most evident threat posed by subsidence, so concrete methods to measure it and thresholds to prevent it have been established (6 mm y^-1^ for the saltmarsh area, National Project Decree 2005). Other threats posed by subsidence to “natural values” would also warrant the adjustment of gas production (National Project Decree 2005), but in those cases three problems might hinder the effective implementation of the “hand on the tap” principle. First, the lack of definition of what the Wadden Sea’s natural values are. We propose that the variables measured here (sediment composition, biomass and benthic community structure) can be considered natural values of the Wadden Sea as they constitute essential components of the ecosystem’s function. Sediment composition and the benthic community might not be the charismatic reason why the Wadden Sea is protected, but these elements are two of the pillars sustaining the ecosystem as we know it today. The second problem is the lack of guidelines about what constitutes an effect. Here, we showed that effects on sediment and community composition might occur, but the lack of measures from before the beginning of gas extraction precludes us from attributing causality. This does not mean that effects do not exist, but rather that it is not possible to unambiguously show them. To do proper monitoring of potential effects, BACI studies are essential. Therefore, we call for the enforcement of properly designed monitoring of the natural values of the Wadden Sea before any activity starts. In cases such as the one studied here, in which before measurements cannot be obtained anymore, we propose that the methods used in this study could be implemented as the standard for identifying potential effects. In cases in which it is methodologically impossible to attribute causation, and given that the Wadden Sea is a protected area, the weight of proof should be reversed, and those exploiting the area should prove no effects are in place. Moreover, by law, the burden of proof is already on the companies exploiting the natural resources showing that their activities have no effect on the Wadden Sea’s natural values (e.g. see Eco Advocacy CLG v An Bord Pleanala. 2023). The third problem that we identify is the lack of a threshold for these effects that, from an ecological perspective, would warrant management consequences. Does the 1 µm y^-1^ decrease in median grain size warrant management consequences? Could this be considered a threat to the natural values of the Wadden Sea? If we do a stringent interpretation of the decree, any effect is reason to reduce or stop drilling.

In conclusion, for a successful implementation of the “hand on the tap” principle in a protected area like the Wadden Sea, it is necessary to clearly define beforehand the relevant variables that represent the natural values, implement proper monitoring for impact assessment, and define threshold values above which effects are not acceptable, otherwise the water will continue to run regardless of where the hand is.

## Author contribution

Allert Bijleveld and Geert Aarts conceived the idea. Allert Bijleveld, Geert Aarts and Paula de la Barra designed the methodology. Paula de la Barra analyzed the data; interpretation of results was discussed by all three authors. Paula de la Barra led the writing of the manuscript. All authors contributed critically to the drafts and gave final approval for publication.

## Acknowledgements

We are very grateful to the current SIBES core-team for collecting and processing the thousands of samples. In alphabetical order they are Leo Boogert, Thomas de Brabander, Anne Dekinga, Job ten Horn, Loran Kleine Schaars, Jeroen Kooijman, Anita Koolhaas, Franka Lotze, Simone Miguel, Luc de Monte, Dennis Mosk, Amin Niamir, Dana Nolte, Dorien Oude Luttikhuis, Bianka Rasch, Reyhane Roohi, Juan Schiaffi, Marten Tacoma, Eveline van Weerlee and Bas de Wit. We also thank all former and current employees and the many volunteers and students who have ensured that the SIBES program has continued in recent years. The RV Navicula was essential for collecting the samples and particularly we thank the current crew Wim Jan Boon, Klaas Jan Daalder, Bram Fey, Hendrik Jan Lokhorst and Hein de Vries. SIBES is currently financed by the Nederlandsche Aardolie Maatschappij NAM, Rijkswaterstaat RWS and the Royal NIOZ.

## Data availability statement

Data will be available at NIOZ Data Archive System upon publication.

