## Supplementary material for "Gas extraction under intertidal mudflats is associated with declines in sediment grain size and minor changes in macrozoobenthic community composition"

Electronic Supplementary Material

**ESM 1: samples per sampling stations**

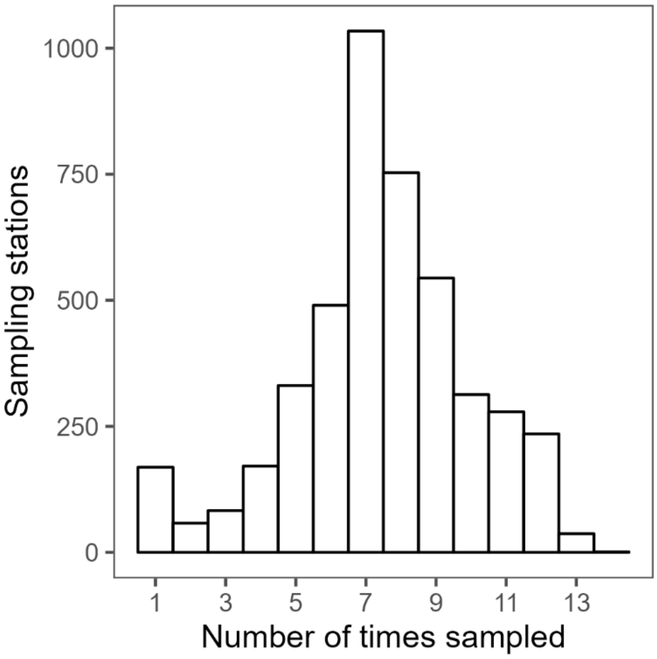


During the synoptic intertidal benthic survey (SIBES), every year we aim to visit and process 4511 sampling stations. However, this is not always possible, so not every station has been visited 13 times between 2088 and 2020. This histogram shows the distribution of samples taken per sampling station during the study period.

**ESM 2: Model results excluding *Arenicola marina***

**Table S1**: Model results for sediment median grain size when excluding *A. marina* as a covariate. Elevation has a nonlinear effect, therefore, three parameter estimates are associated to it.

| Variable | Parameter estimate (mean) | 95% confidence interval |
|---|---|---|
| Subsidence | -1.59 | -3.42 - 0.23 |

| | | |
|---|---|---|
| Year | 0.08 | -0.54 - 0.7 |
| Subsidence : Year | -0.84 | -1.24 – -0.44 |
| Bivalve bank | -9.23 | -9.96 – -8.5 |
| Elevation 1 | -1.22 | -2.15 – -0.29 |
| Elevation 2 | -4.28 | -6.03 – -2.53 |
| Elevation 3 | 4.61 | 3.67 – 5.55 |
| Sampling method | 1.01 | -0.05 – 2.07 |
| Shear stress | 1.42 | 0.89 – 1.94 |
| Slope | -0.01 | -0.27 – 0.26 |
| Topographic position index | 1.45 | 0.45 – 2.45 |

**Table S2**: Model results for mud fraction when excluding *A. marina* as a covariate.

| **Variable** | **Parameter estimate (mean)** | **95% confidence interval** |
|---|---|---|
| Subsidence | 0.39 | 0.22 – 0.55 |
| Year | 0.00 | -0.02 – 0.01 |
| Subsidence : Year | 0.00 | -0.01 – 0.02 |
| Bivalve bank | 0.43 | 0.41 – 0.46 |
| Elevation | -0.11 | -0.13 – -0.1 |
| Sampling method | -0.06 | -0.11 – -0.02 |
| Shear stress | -0.08 | -0.1 – -0.06 |
| Slope | -0.05 | -0.06 – -0.04 |
| Topographic position index | -0.36 | -0.4 – -0.32 |

### ESM 3: SPDE specifications and results

To account for spatial autocorrelation between observations that cannot be explained by the available covariates, a stochastic partial differential equation (SPDE, Lindgren et al., 2011) model was used. We established a non-convex hull around all sampling stations using function 'inla.nonconvex.hull' in R package INLA (Lindgren & Rue, 2015). Within it, we constructed a mesh of triangles with a maximum edge length of 5 km and a cutoff of 1 km (Zuur et al., 2019). We also established a buffer zone around the convex hull with triangles of 25 km maximum edge (Zuur et al., 2019). In this way  a mesh of 1322 nodes was constructed to be used in our models.

**SPDE for sediment median grain size**

Based on DIC values, the best temporal structure for the random field was "AR1" meaning that there was between-year autocorrelation in the median grain size observed at the sampling stations (Table

S3). The the standard deviation of the spatial latent field was 28.27, and the range was 4.1 km, meaning that spatial autocorrelation was negligible at distances larger than that (Fig S1). 95% of the values of this spatial random field fell between -55 and 55 µm.

**Table S3:** Selection between median grain size models with different temporal correlation

| Correlation structure | ΔDIC |
|---|---|
| AR1 | 0 |
| No correlation | 1192 |
| Replicated | 5605 |

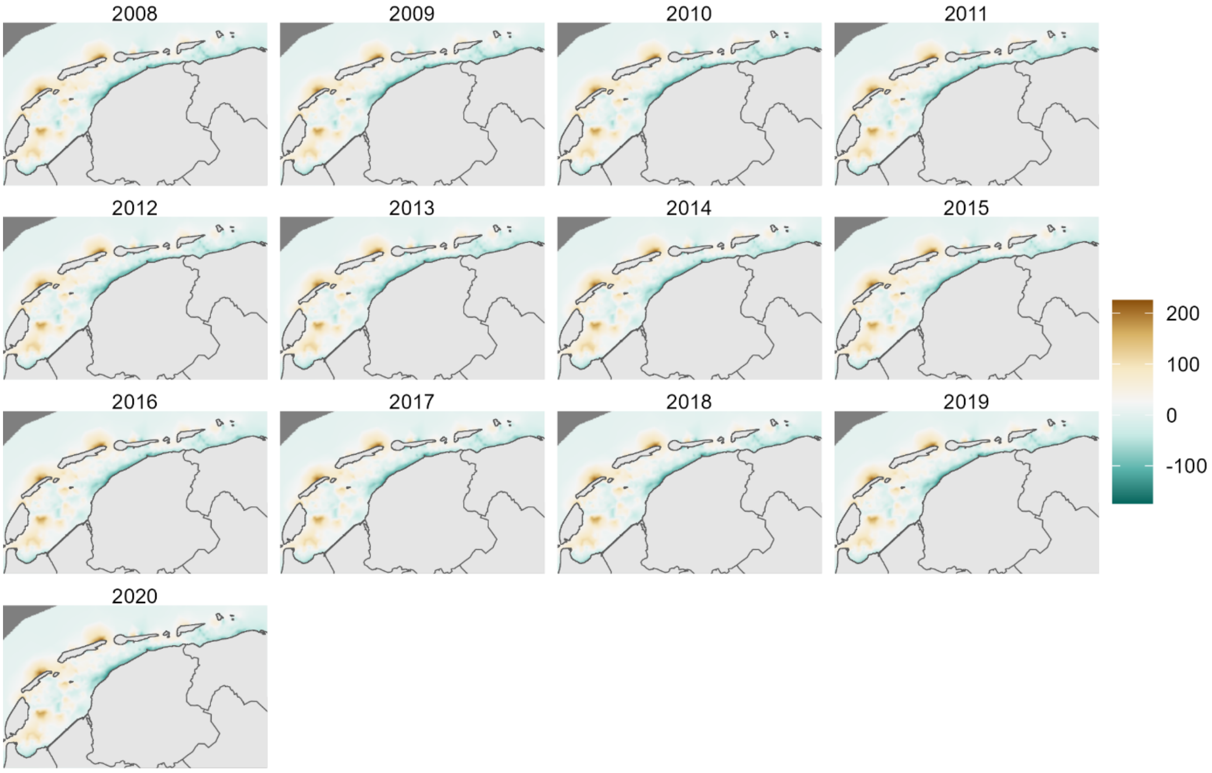


**Fig S1:** Random fields for the sediment median grain size model.

**SPDE for mud fraction**

The best temporal structure for the random field was "AR1" (Table S4). The random field had a standard deviation of 1.19 and a range of 8.7 km, and 95% of the values fell within 0.09 and 0.69 (Fig S2).

**Table S4:** Selection between mud fractions models with different temporal correlation

| Correlation structure | ΔDIC |
|---|---:|
| AR1 | 0 |
| No correlation | 1168 |
| Replicated | 5046 |

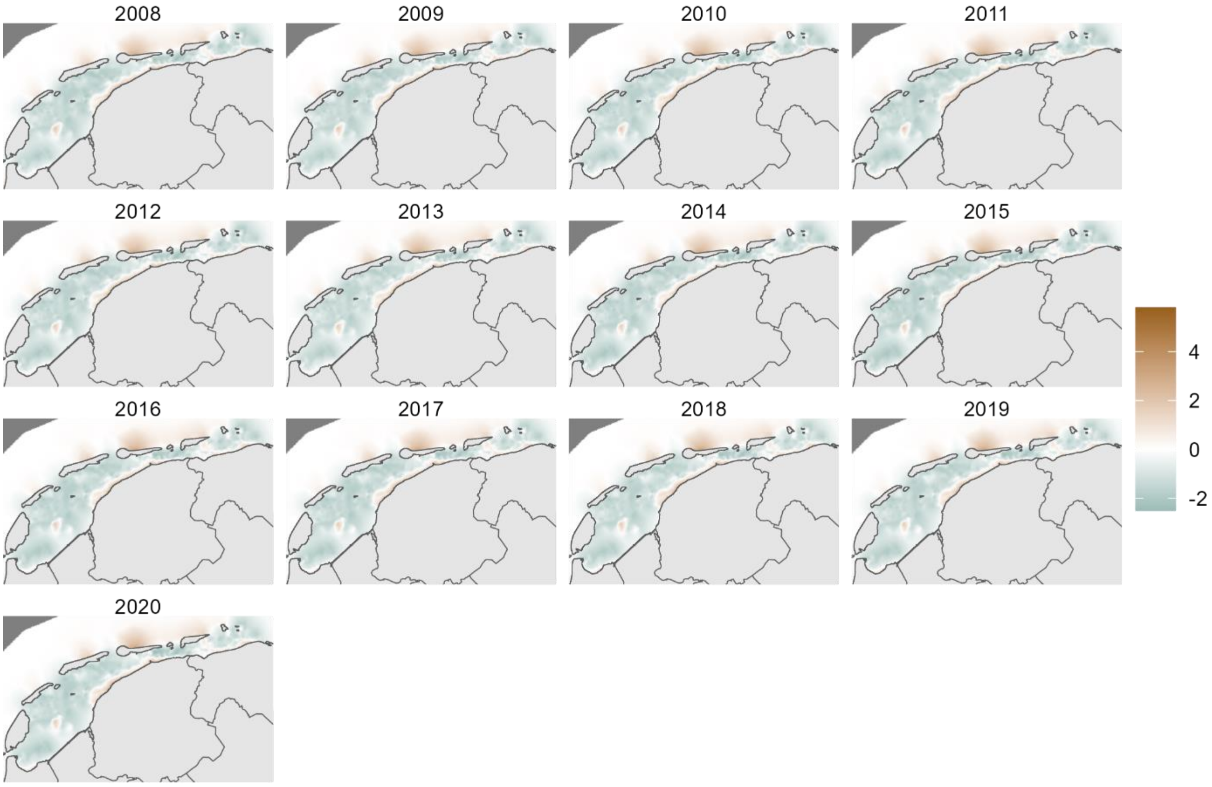


**Fig S2:** Random fields for the mud fraction model.

**SPDE for total biomass**

The best temporal structure for the random field was "AR1" meaning that there was autocorrelation between the random components of each year (Table S5). The standard deviation of the random field was 1.13 with 95% of the values between 0.25 and 4.35 g $m^{-2}$. The range of spatial autocorrelation in the random field was 2.15 km (Fig S3).

**Table S5:** Selection between models with different temporal correlation

| Correlation structure | ΔDIC |
|---|---|
| AR1 | 0 |
| Replicated | 1582 |
| No correlation | 12638 |

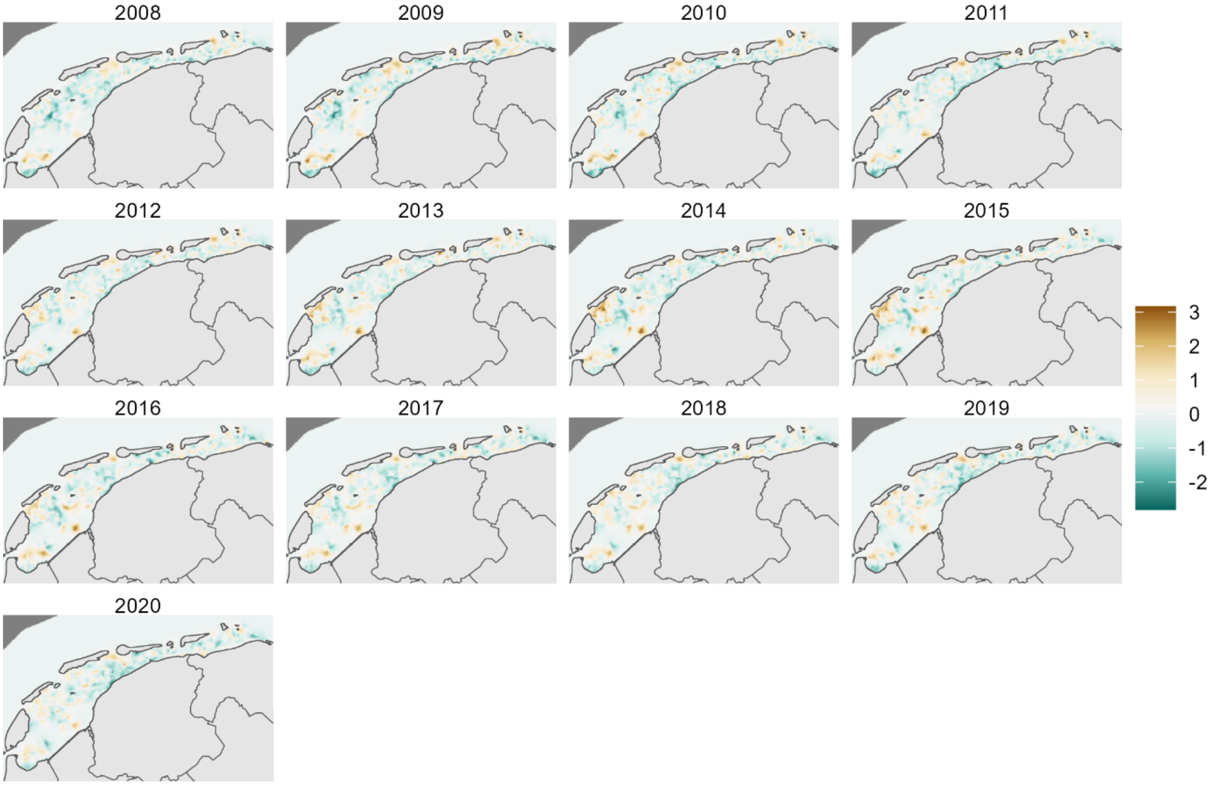


**Fig S3:** Random fields for the biomass model.

**Interpretation of SPDE**

While several environmental variables significantly influence the observed variation in the response variables, the random spatial field still explains a large part of the variability in all three models. For example, for the median grain size model, the random field can reach values greater than 55 µm (Fig S1), while average median grain size is 154 µm, meaning that for some areas, the unexplained variation is large, and the random component is an important part of the prediction. In addition, for median grain size and mud fraction, the annual random field estimates are also biased towards negative values, while we would expect them to be zero on average (Fig S4). This observed bias of the random field might be absorbing some of the annual variation that could not be captured by the continuous variable 'year', nor by any other environmental variables included in the model.

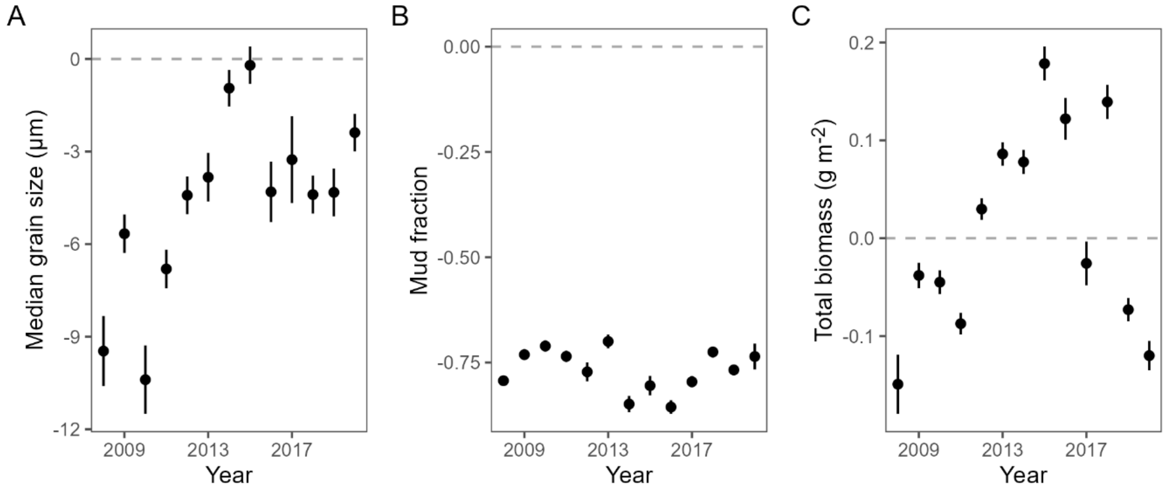


**Fig S4**: Values that the random field takes at the location of the stations (mean ± se). The plots are on the link scale.

### ESM 4: "Fake subsidence" controls

We established "fake subsidence areas" (Fig S5) and compared their median grain size and mud fraction with that of the rest of the Wadden Sea excluding the real subsidence area. We aimed to select one area per tidal basin that included a similar number of samples as the subsidence area, and that did not include areas affected by other sources of land subsidence (Fokker et al., 2018; Hoeksema et al., 2004). We allowed "fake subsidence aeras" to spill out of the tidal basins' limits, but still we did not always achieve to delineate areas with high enough number of samples. In the end, we were able to established five "fake subsidence areas" (Fig S5, Table S6).

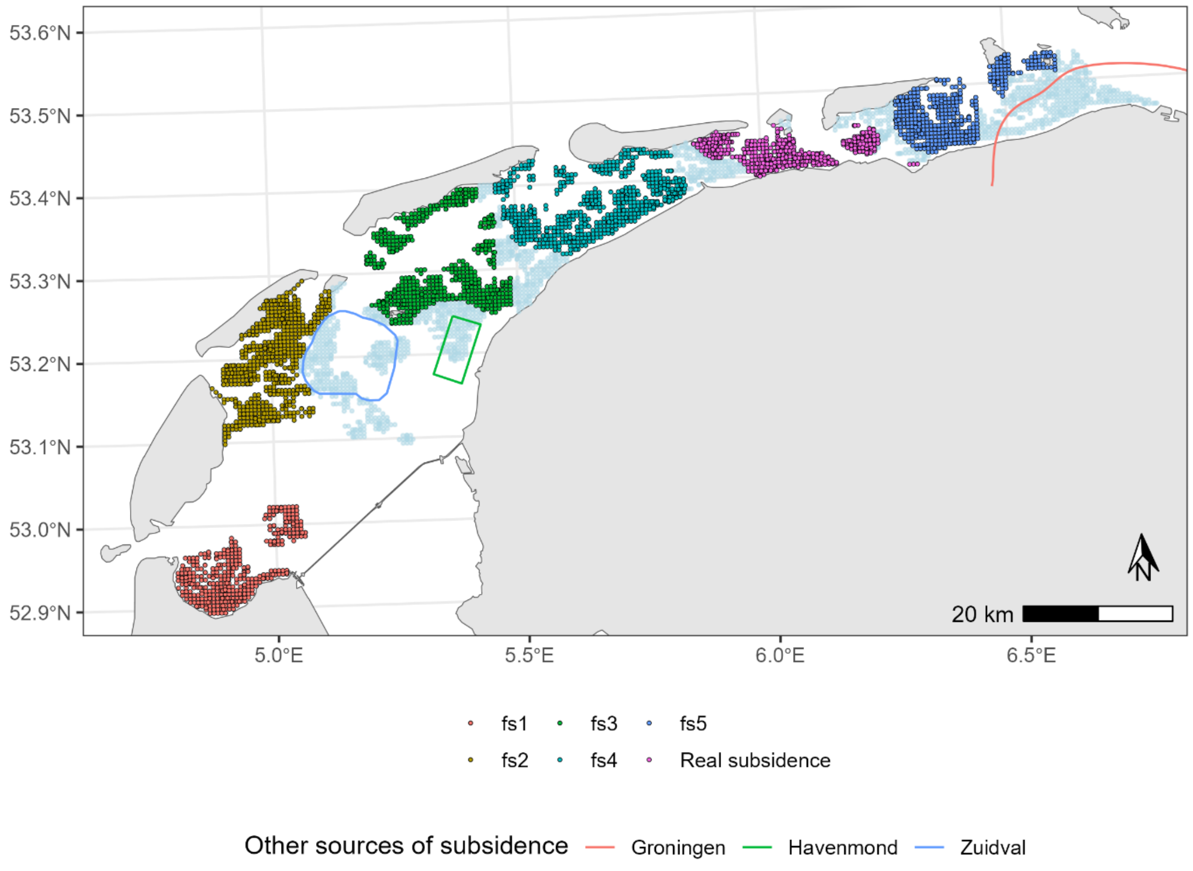


**Fig S5:** Samples conforming the five fake subsidence (fs) areas, the samples from the real subsidence area, and contours for other sources of subsidence. Points in light blue show the rest of the sampling stations. Contour in blue shows the area affected by subsidence resulting from gas extraction from the Zuidwal field (Hoeksema et al., 2004), contour in green shows the area licensed for salt mining (Fokker et al., 2018 no contours for subsidence are available for this field). Contour in red shows subsidence associated to gas extraction from the Groningen field (Operationele Strategie voor Gasjaar 2022-2023).

We ran five models for each response variables (median grain size, mud fraction and total biomass), comparing each of the fake subsidence areas with the rest of the Wadden Sea minus the real subsidence area. We used the same covariates as in the original model. We aimed at using the same

SPDE specifications too, but for some models we needed to adjust the priors to achieve a good fit (median grain size: fake subsidence 2 and 4).

We did not find significant differences between "fake subsidence" areas and the rest of the Wadden Sea neither for median grain size nor for mud fraction (Table S6, Fig S6), but we did find differences for the biomass model in two "fake subsidence" areas relative to the rest of the Wadden Sea (Table S6, Fig S6 C).

**Table S6:** Number of samples and sampling stations within the real and different fake subsidence areas, and main results for the median grain size and mud fraction models. Ranges between brackets are 95% confidence intervals

| | Fake subsidence 1 | Fake subsidence 2 | Fake subsidence 3 | Fake subsidence 4 | Fake subsidence 5 | Real subsidence |
|---|---|---|---|---|---|---|
| Number of samples | 3077 | 4032 | 4104 | 4109 | 3223 | 3866 |
| Number of sampling stations | 384 | 582 | 543 | 619 | 410 | 488 |
| **Median grain size model** | | | | | | |
| Subsidence parameter | 0.00 (-1.96 – 1.96) | 0.47 (-1.41 – 2.35) | 0.24 (-1.57 – 2.05) | 0.47 (-1.41 – 2.35) | 1.00 (-0.85 – 2.84) | -1.74 (-3.58 – 0.1) |
| Subsidence : Year parameter | 0.05 (-1.41 – 1.52) | 0.57 (0.02 – 1.11) | -0.08 (-0.46 – 0.31) | 0.57 (0.02 – 1.11) | 0.4 (-0.01 – 0.81) | -1.13 (-1.56 – -0.69) |
| **Mud fraction models** | | | | | | |
| Subsidence parameter | 0.01 (-0.85 – 0.88) | -0.16 (-0.38 – 0.07) | 0.06 (-0.08 – 0.21) | -0.11 (-0.3 – 0.08) | 0.10 (-0.06 – 0.26) | -1.74 (-3.58 – 0.1) |
| Subsidence : Year parameter | -0.02 (-0.06 – 0.02) | 0.00 (-0.02 0.02) | -0.01 (-0.02 – 0.01) | -0.01 (-0.03 – 0.01) | 0 .00 (-0.02 – 0.02) | -1.13 (-1.56 – -0.69) |
| **Total biomass model** | | | | | | |
| Subsidence parameter | 0.00 (-0.05 – 0.04) | 0.03 (0 – 0.06) | 0.02 (-0.01 – 0.05) | -0.03 (-0.06 – 0.01) | 0.01 (-0.02 – 0.05) | -0.03 (-0.07 – 0.01) |
| Subsidence : Year parameter | -0.02 (-0.43 – 0.39) | **-0.48 (-0.7 – -0.25)** | **-0.26 (-0.5 – -0.03)** | -0.09 (-0.36 – 0.18) | -0.32 (-0.59 – -0.04) | 0.18 (-0.16 – 0.52) |

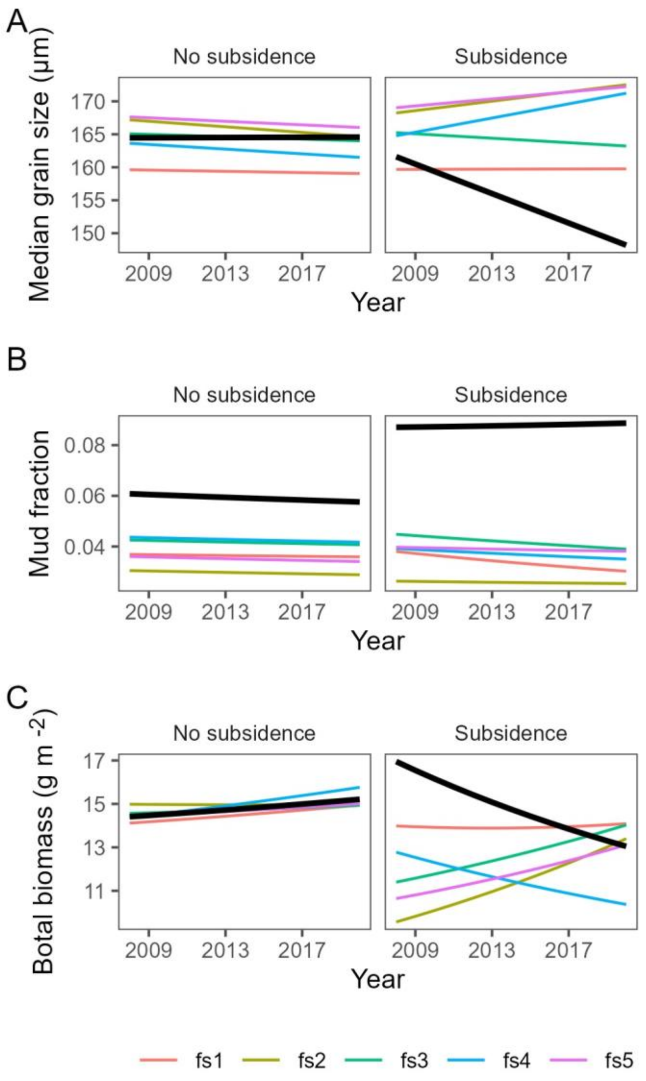


**Fig S6**: Predictions for the models using fake (fs, colored thin lines) and real subsidence areas (thick black lines). **A**. Predictions for median grain size. **B**. Predictions for mud fraction. **C**. Predictions for total biomass.

**Species-specific analysis**

Similarly, we ran additional analysis for the species specific analysis, using the biomass of species within the "fake subsidence" areas to calculate the log biomass ratio between these and the rest of the Wadden Sea for each year.

We found that the ‘Elevation use : Year’ interaction parameter was different from zero in one of the controls (Table S7).

**Table S7:** Main results for the log biomass ratio models using “fake” and real subsidence areas. Ranges between brackets are 95% confidence intervals and are shown in bold when they exclude zero.

| | Fake subsidence 1 | Fake subsidence 2 | Fake subsidence 3 | Fake subsidence 4 | Fake subsidence 5 | Real subsidence |
|---|---|---|---|---|---|---|
| **Population trends** | | | | | | |
| Year parameter | 0.01 (-0.00 – 0.02) | 0.01 (-0.01 – 0.02) | 0.00 (-0.01 – 0.01) | -0.00 (-0.01 – 0.01) | -0.00 (-0.01 – 0.01) | 0.00 (-0.00 – 0.01) |
| Elevation use : Year parameter | -0.01 (-0.02 – 0.00) | 0.01 (-0.01 – 0.02) | 0.01 (-0.00 – 0.02) | -0.02 **(-0.03 – -0.01)** | -0.00 (-0.01 – 0.01) | -0.01 **(-0.02 – -0.00)** |
| ΔAIC (null model – full model) | 0.81 | 1.78 | -3.82 | 9.44 | 0.83 | 3.96 |

### ESM 5: selection of species for the analysis of species-specific effects of land subsidence

We wanted to include in this analysis as much species as possible to identify effects in common and rare species. However, for species with very low frequency of occurrence, the estimated average biomass would have high uncertainty and might skew the analysis. For this reason, we studied how the ratio between species biomass’ mean and standard deviation changes depending on the species’ frequency of occurrence. We plotted this relationship (Fig S7) and established the inclusion threshold considering a compromise between the stabilization of the ratio between species biomass’ mean and standard deviation, and the number of species above this threshold. As a result, only those species that occurred in at least 100 samples each year (~2% of samples) were included in this analysis.

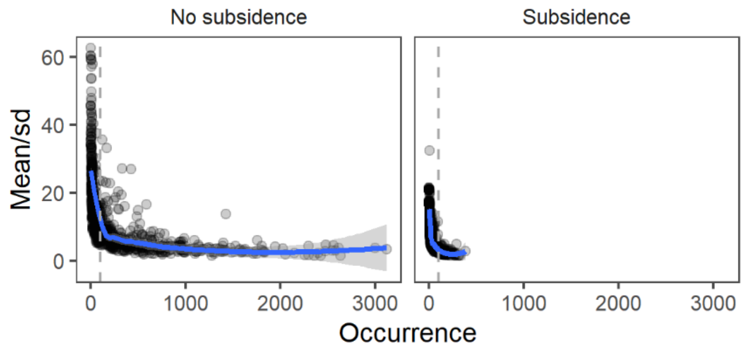


**Fig S7:** Changes in the ratio between mean biomass and standard deviation (sd) within and outside the subsidence area, depending on frequency of occurrence. Means and sd were calculated on a yearly basis for each species.

### <u>ESM 6: Nonlinear relationships</u>

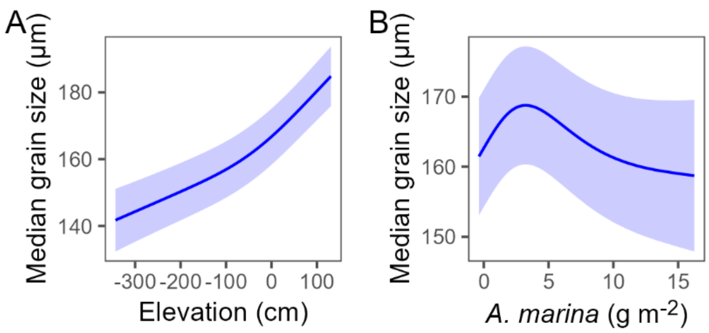


**Fig S8:** Smoothers used in the median grain size model for (**A**) Elevation and (**B**) *Arenicola marina*.

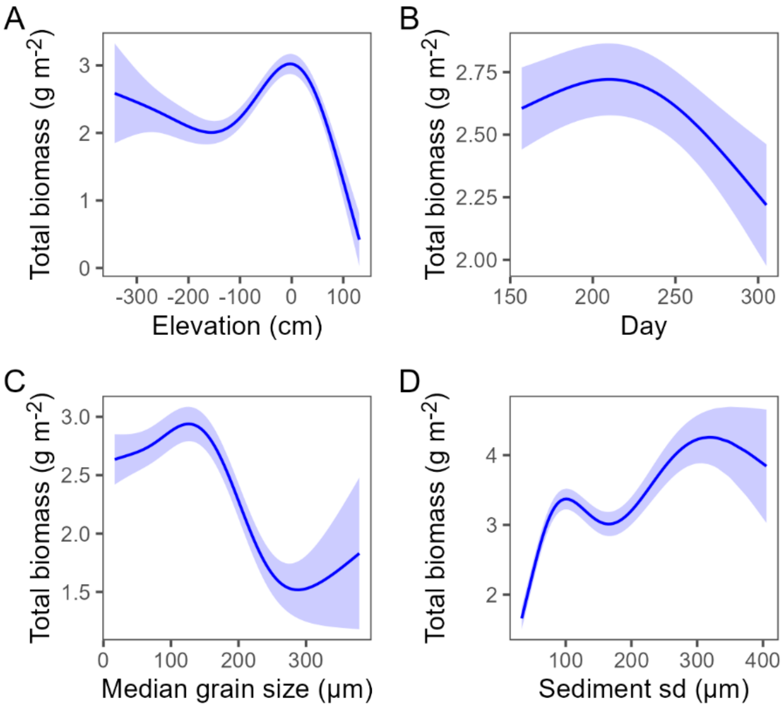


**Fig S9:** Smoothers used in the total biomass model for (**A**) Elevation, (**B**) Day of the year, (**C**) Sediment median grain size and (**D**) Sediment grain size standard deviation.
